# Comparative experimental studies on Trypanosoma isolates in goats and response to diminazene aceturate and isometamidium chloride treatment

**DOI:** 10.1101/2023.02.26.530157

**Authors:** Abrham Ayele, Shimelis Dagnachew

## Abstract

The objective of this study was to characterize and compare the clinically visible pathological features and drug resistance patterns of two Trypanosoma isolates from two tsetse infested areas of northwest Ethiopia in experimentally infected goats. From the 37 trypanosome free goats, two goats (trypanosome donors) were used to take trypanosome isolates from naturally infected cattle (one from Jawi and the other from Jabitehenen areas). The remaining thirty five goats were randomly assigned into seven experimental groups, each containing five goats. These groups were again randomly selected, and three of them—Group 1 (JWI-1), Group 2 (JWI-DA), and Group 3 (JWI-ISM)—were inoculated with the Trypanosoma of Jawi isolate. Group-4 (JBI-1), Group-5 (JBI-DA), and Group-6 (JBI-ISM) were inoculated with the Trypanosoma of Jabitehenan isolate. The remaining group, Group 7 (NIC), was under negative control. Each experimental goat received 2 ml of trypanosoma positive blood at the 1×10^6^ parasites/ml from donor goats through the jugular vein. Group NIC received 2 mL of sterile water as a negative control. For ten weeks following infection, parameters such as parasitaemia, body weight, PCV, and hemoglobin value were measured once per week. When peak parasitaemia was detected on day 14 of post infection, trypanocidal treatment was administered. Diminazine diaceturate (DA) was given at a dose of 28 mg/kg, and isomethamedium chloride (ISM) was given at a dose of 4 mg/kg. Trypanosomosis was detected on days 5 and 6 of post-infection (Pi) in Jabitehenan and Jawi, *T. congolense* isolates from infected groups, respectively. When peak parasitaemia was detected on day 14 of post infection, trypanocidal treatment was administered. 7 mg/kg for Group 2 and Group 5 and 1 mg/kg for Group 3 and Group 6 were the treatment doses for DA and ISM, respectively. All of the infected groups (groups 1 and 4; positive control) had severe clinical signs and recumbence within 27-59 days of infection. The mean PCV, total RBC, and Hgb concentration values of Jabitehenan isolate infected groups were significantly (*P*<0.05) lower than Jawi isolate infected groups, and severe clinical signs in group 4 occurred earlier than in group 1. These findings point to the presence of inter-isolate *T. congolense* variation. Neither of the DA nor ISM treatment groups attained complete recovery, and this shows the presence of disease resistance. A molecular-based study was this study’s limitation, and it is recommended to confirm interisolate differences.

## 1. INTRODUCTION

Trypanosomes are parasitic protists that circulate in the bloodstream and tissue fluids of their mammalian hosts (1). *Trypanosoma brucei* spp., *Trypanosoma congolense, Trypanosoma vivax, Trypanosoma evansi* and *Trypanosoma equiperdum* and *Trypanosoma cruzi* are regarded as the salivarian groups (2) following their character of growing in the tsetse fore or mid gut so that transmission is via tsetse biting. In contrasting to this character, those trypanosoma spp. growing in hind guts of their arthropod vectors (like in triatomid kissing bug) are called as Stercorarian trypanosome. For the salivarian pathogenic trypanosomes the subgenera include Trypanozoon, Duttonella, Nannomonas, and Pycnomonas, of which the first three account for the vast majority of human and animal infections (3).

The salivarian trypanosomes belonging to the subgenus Nannomonas (*T. congolense* and *T. simiae*) are major pathogens of livestock in sub-Saharan Africa (4). In Africa, the Americas and Asia, these diseases, which in some cases affect humans, result in significant illness in animals and cause major economic losses in livestock (5).

The Abay (Blue Nile) basin of Northwest Ethiopia is among the risk areas victims to this parasite (6). In sub-Saharan Africa, *Trypanosoma congolense* is the most pathogenic trypanosome species infecting livestock. Isoenzymatic differences in different *T. congolense* isolates resulted in a subdivision of the *T. congolense* species in to several types; the *T. congolense* savannah type, *T. congolense* Tsavo type, *T. congolense* forest type and the *T. congolense* Kilifi type (7; 8). Geysen *et al*. (9) tried to analyse this parasite’s sub-species based on the PCR-RFLP approach targeting the 18S small ribosomal subunit gene and classified it into three different types; the savannah, forest and the Kilifi types. Each subgroup possesses a unique satellite DNA sequence (10). The ability to identify these subgroups accurately opens the way for systematic studies of their host range, distribution and pathogenicity. The pathogenicity appears to vary depending on which type or strain of *T. congolense* is involved for infection (12). For example *T. congolense* savannah type is found to be the most pathogenic to mice and cattle (12).

Although the mechanisms involved are poorly understood, weight loss, anemia, enlargements of lymph nodes and immune-suppression are taken as pathological symptoms of trypanosomosis (13). Following these unspecific clinical signs, it has been difficult to differentiate it from other infections and to control it especially in remote areas of Ethiopia where there is no accesses laboratory diagnosis.

As chemotherapy is the most widely used and more effective way of trypanosomosis control, similar kinds of trypanocidal drugs have been used for more than 60 years in Ethiopia (14). According to Assefa *et al*. (15) treatments and prevention of African animal trypanosomosis has been relied essentially on three drugs namely: Homidium chloride/homidium bromide, diminazene aceturate and isometamidium chloride (14).However, resistance to one or more of the three-trypanocidal drugs used in cattle has been reported(16). The compounds of diminazene and isometamidium have been termed as a sanative pair for the control of bovine trypanosomosis (17). However, reports are still documented for the expression of multiple drug resistance phenotypes (18;19;20;21)

Since the three types of *T. congolense* were acknowledged, the varieties of pathogenicity between strains become a concern (12). However, there is limited information about *T*.*congolense* strain type, pathogenesis and drug resistant patterns in Ethiopia, particularly in the northwest region. Information on the impact of *T. congolense* infections on hematological values is also limited too. Therefore, the present study determined and compared clinical findings, hematological values and drug resistance patterns induced by two *T. congolense* isolates of northwest Ethiopia.

## 2. MATERIALS AND METHODS

### 2.1. Experimental animals purchasing and their managements

Thirty seven apparently healthy indigenous goats with relatively the same age (between 13-15 months) and the same weight (16-18kg) were purchased in Gondar town where it is considered as free of tsetse fly and trypanosomosis. Then they transferred to the university of Gondar experimental animals’ premises. Goats were acclimatized to the new environment handling and feeding conditions for one month prior to the beginning of the experiment. Throughout the acclimatization period, animals feeding, drinking, and other activities were observed daily. Goats’ weight, packed cell volume (PCV), and rectal temperature were measured weekly.

Faecal samples of each experimental goat were diagnosed with gross faecal examination, sedimentation and floatation techniques weekly too. All animals were also dewormed with ivermectin (1% injection, Nor-brook, UK) and albendazole (300mg) (Albenda-QK, Chengdu Qiankun, China) in 2 weeks interval.

During the experimental period, animals were fed *ad libitum* on hay, Green elephant grass and water. Supplements of concentrates and mineral licks were given two times per day at around 8:00 am (morning) and dusk (5:00 pm). Any experimental animals recommended for postmortem examination and those animals with severe clinical manifestations with PCV below 15 % were humanly euthanized using jugular vein injection of an overdose of sodium Phenobarbital (13). After completion of the postmortem examination, animals’ carcass were burned deep in to the ground and buried in it. The University of Gondar Animal Research Ethics Review Committee was also authorized this research for its safe and sound experimental animal management.

### 2.2. Field parasites isolation and transportation to experimental site

Trypanosome parasite were isolated from cattle (oxen) of the two tsetse infested areas of northwest Ethiopia: one from Jabitehenan district (JBI-D1=Jabitehenan donor 1), located at 10^°^42’N and 37^°^16’E with an altitude range of 1500-2300meter above sea level (m.a.s.l.) and the second were from Jawi district (JWI-D1 = Jawi donor 1), Located at 11^°^36’N and 37^°^23’E with an altitude range from 648-1300 m.a.s.1 (Fig.1). Both areas comprise a variety of different vegetation types including savannah, woodland, riverine and cultivated lands. Permanent and seasonal rivers also made the areas ideal for tsetse fly habitation.

**Figure 1:**
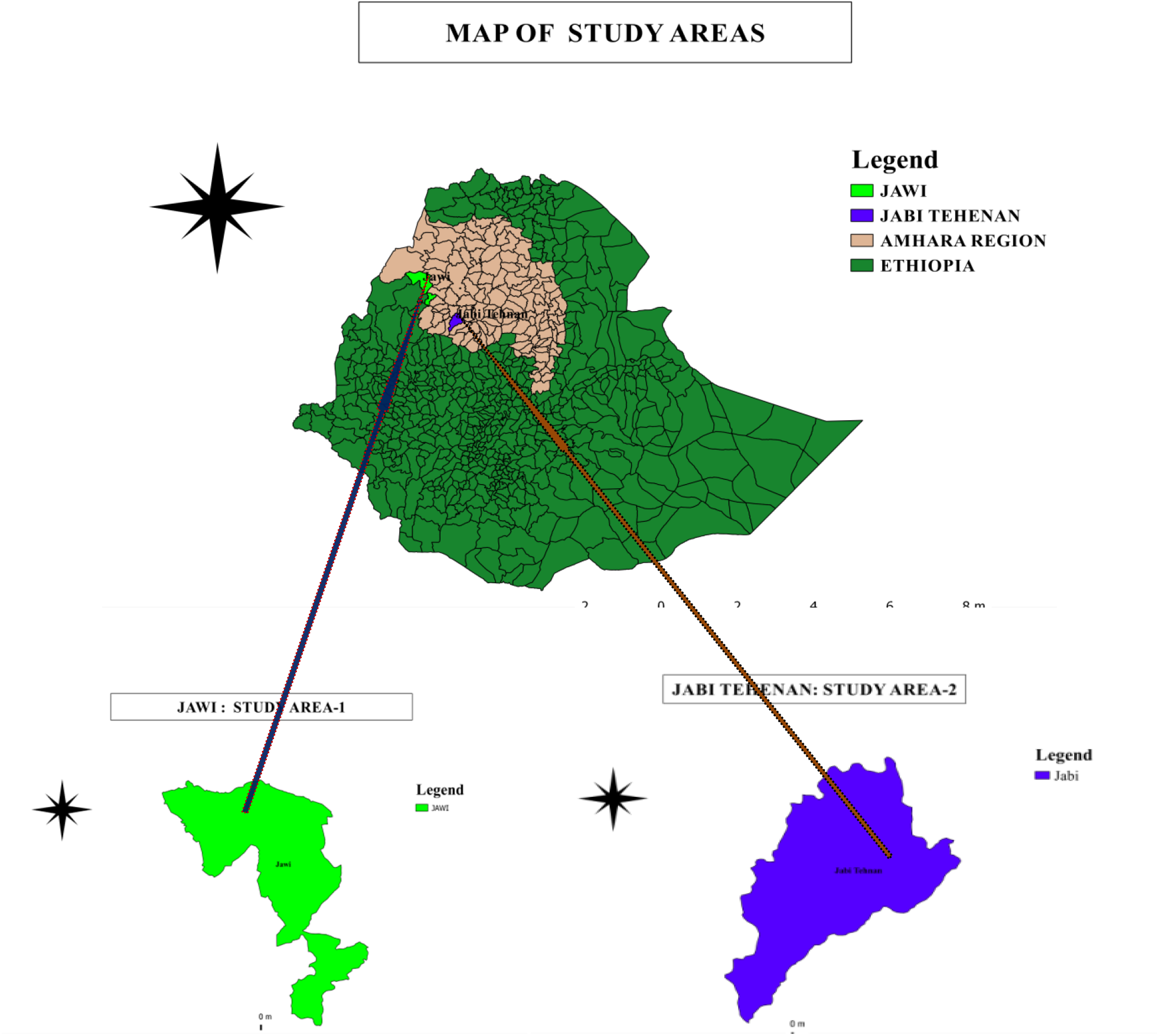
Map showing *Trypanosoma congolense* isolated areas used for the experiment. From Jabitehenan region, a six years old ox used for tracking and managed in free range system was found to be infected with *T*.*congolense*. Its parasitaemia level was averagely 2 parasites per field of microscope slide. 2ml of blood was drained from jugular vein of that ox and given to a healthy male goat that was acclimatized for one month at the experimental site and taken to field for trypanosome transportation purpose.

Similarly, a six years old ox used for tracking and managed in free range system was also found to be infected with *T*.*congolense* in Jawi area with similar parasitaemia level as found in Jabitehenan district. 2ml blood from jugular vein was drained from that ox and given to a healthy male goat through its jugular vein. These donor goats were then returned to experimental sites present at the University of Gondar, located at 12°45’N and 37°45’E with an altitude of 2133m.a.s.l.

The isolated parasites were confirmed as *T. congolense* by observing the parasite’s movement and morphology with wet blood film, buffy coat and thin smear followed by Giemsa staining technique examinations (22). *T. congolense* was identified morphologically based on the absence of free flagellum and marginal and medium sized kinetoplasts. Its movement is vibrating on its position while one of its extremities is attached to the blood cell (22). ***However, the absence to molecular test to identify T. congolense sp. is the limitation of this study***.

### 2.3. Experimental groups and trypanosome challenges

As two goats are trypanosome donors, the remaining thirty five experimental goats were randomly divided into seven groups of five goats in each group (Table1). Randomization was done by lottery method using the list of their identification code in each experimental animal’s ear tag. Then groups are randomized again to be infected with Jawi or Jabitehenan isolates. Group 1 (JWI-1) were infected with *T. congolense* isolated from Jawi area and left as infected control; Group 2 (JWI-DA) were infected with *T. congolense* isolated from Jawi area and then treated with diminazine di aceturate (DA); Group-3 (JWI-ISM) were infected with *T. congolense* isolated from Jawi area and latter treated with isomethamidium chloride (ISM); Group-4 (JBI-1) were infected with *T. congolense* isolated from Jabitehenan area and left as infected control; Group-5 (JBI-DA) were infected with *T. congolense* isolated from Jabitehenan area and then treated by DA; Group-6 (JBI-ISM) were infected with *T. congolense* isolated from Jabitehenan district and followed by ISM treatment and Group-7(NIC) were the non-infected control group. All anti parasitic treatments were done by animal physician carefully at peak parasitaemia level (table 1).

**Table 1:** Experimental groups of goats challenged with *T. congolense* isolates from Jawi and Jabitehenan areas of northwest Ethiopia.

| Group | Infection status | trypanosome isolates |  | Treatment status |  |
| --- | --- | --- | --- | --- | --- |
|  |  | Jabitehenan isolate | Jawi isolate | DA at 7 mg/kg | ISM at 1 mg/kg |
| **Group-1 (JWI-1)** | Infected |  | X |  |  |
| **Group-2 (JWI-DA)** | Infected |  | X | X |  |
| **Group-3 (JWI-IMM)** | Infected |  | X |  | X |
| **Group-4 (JBI-1)** | Infected | X |  |  |  |
| **Group-5 (JBI-DA)** | Infected | X |  | X |  |
| **Group-6 (JBI-ISM)** | Infected | X |  |  | X |
| **Group-7 (NIC)** | None infected control | - | - | - | - |

The level of parasitaemia in the donor animals was estimated according to the protocol of rapid-matching method described by Herbert and Lumsden (23). Each experimental animal, except group 7 (NIC) received 2 ml of infected blood intravenously from their respective isolates. Inoculated blood was taken from jugular vein of donor goats having a parasitaemia level of 1×10^6^ trypanosomes per ml of blood. The approximate parasitaemia of the infected blood was adjusted using dilution factors.

### 2.4. Clinical and parasitological examinations

Experimentally infected animals were examined daily for their clinical parameters. PCV determination; hemoglobin values and body weight (before they took feed) were measured once per week throughout the study period. Blood samples from each animal’s ear vein were examined daily until the detection of parasites, then weekly using the standard parasitological methods of wet blood smear microscopy and the buffy coat technique (22) to determine the degree of parasitaemia and detection of relapse of the parasite (24). For all parasitological examination, blood samples were aseptically obtained from jugular vein using Ethylene diamine tetra acetic acid (EDTA) coated vacutainer tube and from ear vein directly into heparinized capillary tubes. For each sample at least 2 wet smears and thin smears were made from blood and buffy coat and took the value of the average result to reduced procedural imperfect. After detection of the parasite in the blood of experimental goats, 4 thin blood smears were prepared from a single sample (blood and buffy coat (2 from each)) and stained with Giemsa solution for observation of trypanosome morphology via microscopy as described by Kagira, *et al*. (25).

#### 2.4.1. Haemathological parameters

Packed cell volume (PCV), haemoglobin (Hgb) concentration, red blood cells (RBC) count, erythrocyte indices (mean corpuscular volume (MCV), mean corpuscular haemoglobin (MCH) and mean corpuscular haemoglobin concentration (MCHC)), and total and differential white blood cell (WBC) count were measured weekly from a jugular vein blood sample collected using EDTA coated vacutainer tubes. PCV was measured by the haematocrit centrifugation technique using a Hawksley microhaematocrit reader. Total RBC and WBC counts were carried out manually using the improved Haemocytometer. Hgb concentration was measured by the Sahili’s Acid-Haematin method. The following formulas (26) were used to calculate erythrocyte indices from the above haematological values:

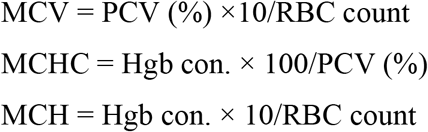

Thin blood smears were stained with Giemsa solution for differential WBC counts, which were based on 100 cells per slide according to their staining reactions; shape of the nucleus, and presence or absence of granules in their cytoplasm (26).

#### 2.4.2. Drug sensitivity tests protocol

The trypanocidal drugs used in the experiment were diminazene aceturate (DA) 2.36g (Veriben made in Libourne France (lot. 827A1 exp.03/2019) and ISM (Isometamedium chloride 125mg (Veridium) manufactured in Libourne-France (Lot 198A1 exp-06/19).

DA was injected for JWI-DA and JBI-DA groups as a 7% solution at a dose of 7 mg/kg body weight and ISM was injected for JWI-ISM and JBI-ISM groups as a 1% solution at a dose of 1 mg/kg of body weight of experimental goats to test the efficacy of high dose of these drugs based on the previous established drug resistance test protocol set by Eisler *et al*., (27). Sterile distilled water was used to dissolve appropriate quantities of the drugs. Drugs, according to the respective group (see table 1), were administered through deep intramuscular route (IM) on the basis of experimental goats body weight after all of them in each group showed peak parasitaemia (1×10^6^ trypanosomes per ml of blood). Since the treatment date, goats were monitored for trypanosome parasitaemia by the wet smear and Buffy coat technique three times a week for 100 consecutive days (27).

#### 2.4.3. Post mortem examination

Experimental animals with sever clinical sign accompanied by recumbence, revealing PCV value less than 15% at any point of the experimental time and all of the infected experimental animals at the end of the experiment were humanly euthanized using high dose Phenobarbital. Two animals from the none infected control groups were also purposively euthanized by the same mechanism for visceral lesion evaluation and comparisons. The postmortem examinations were done cautiously on euthanized animals and each organ was examined carefully to detect the abnormalities. Different lesions from each animal were examined and compared with a grave recording of each of the findings in a data collection sheet.

### 2.5. Data analysis

All data collected were entered into an excel spread sheet and imported into statistical package for social science (SPSS) version 20 statistical software. Differences in haematological values, rectal body temperature and body weight loss measured between groups were assessed by using one-way analysis of variance (ANOVA). Independent t-test was used to compare mean parameters between infections of the two areas *T. congolense* isolates. Paired t-test was used to compare the mean PCV values and body weight of animals before and after treatment. The mean PCV and mean body weight improvement after treatment was calculated using the following formulas (28).

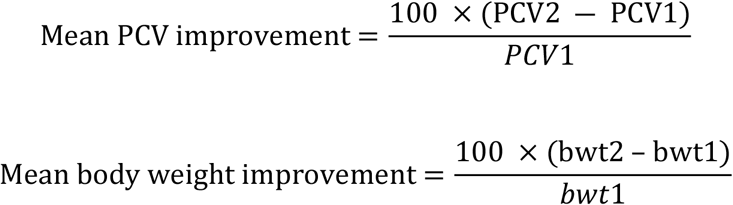

Different tables, graphs and pictures were also used to describe the different experimental values. Post mortem results were recorded carefully and described accordingly. The significant level was set at (*P*< 0.05).

## 3. RESULTS

### 3.1. Development of parasitaemia

Parasitaemia in field infected goats with parasitic cattle blood was detected on day 11 and day 13 of post infection (pi) and reached peak parasitaemia on day 26 and 29 of pi for Jabitehenan and Jawi isolates respectively. In the experimental infected goats parasitaemia was observed on day 5 of pi in Jabitehenan isolate infected groups and on day 6 of pi for Jawi isolate infected groups and it reached at peak load on day 14 of pi in both isolates infected groups Figure (2).

**Figure 2:**
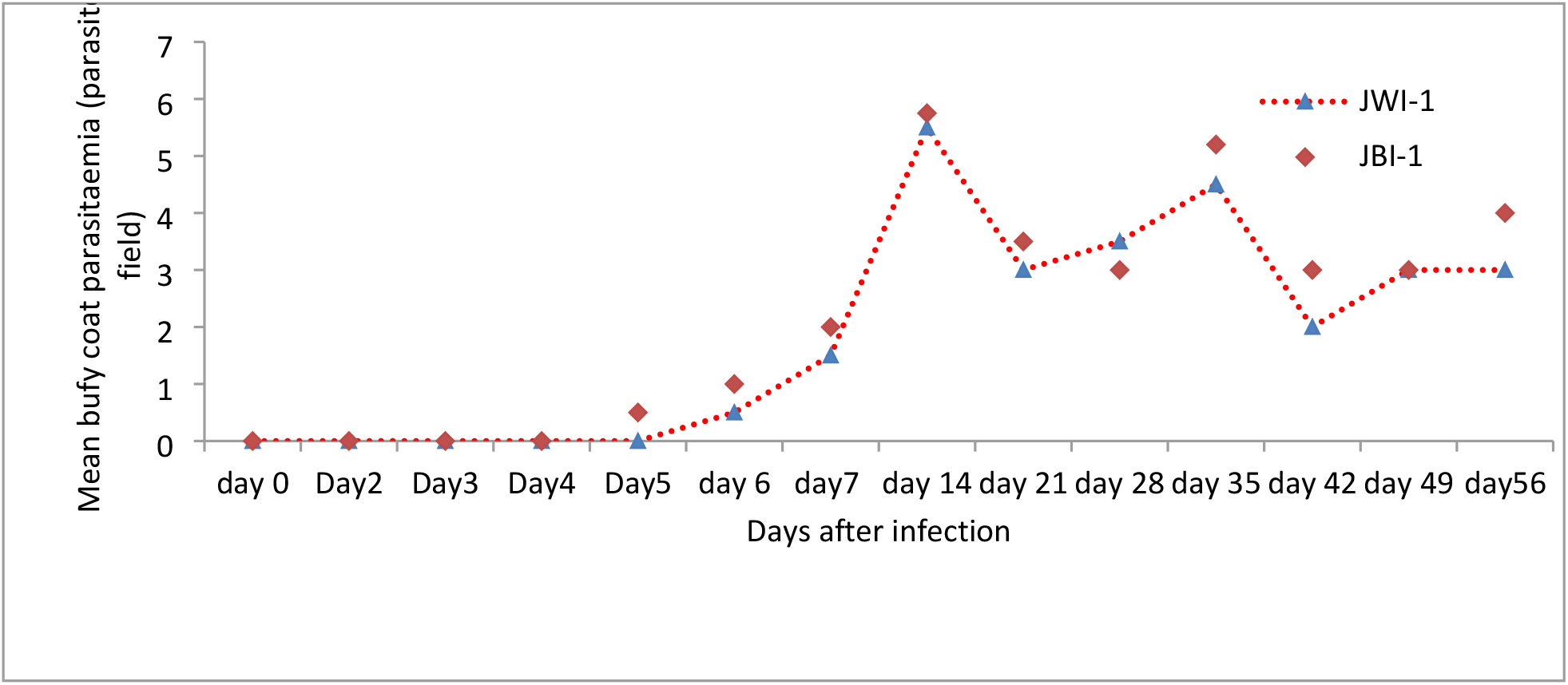
Mean parasitaemia observed during the study period in experimentally infected goats with *T. congolense* isolates from two tsetse infested areas (Jawi (JWI-1) and Jabitehenan (JBI-1)) of northwest Ethiopia.

### 3.2. Clinical findings

Main clinical findings observed in all infected goats were; weakness, fever, rough hair coat, reduced feed intake, enlarged superficial lymph nodes, pallor of the mucus membranes and weight loss (Fig. 3). Diarrhea was observed in two animals of Group JWI-1 and in three animals of Group JBI-1. Sub mandibular edema was observed in three goats’ of JBI-1 group and in a goat from JWI-1 group. Lacrimation was observed in one goat from JWI-1 group and two goats of JBI-1 group. All goats under the infected control groups of both isolates got severe illness within the range day of 27-59 of Pi. However, the time duration for the occurrence of severe clinical signs was short in Groups JBI-1. Three goats infected with Jawi isolates and all goats infected with Jabitehenan isolates of *T*.*congolense* were acquiring recumbence in the second wav e of parasitaemia (day 28-42 pi).The overall mean body temperature and body weight changes observed in the infected and none infected experimental animals during the study period is summarized in Table 2.

**Figure 3:**
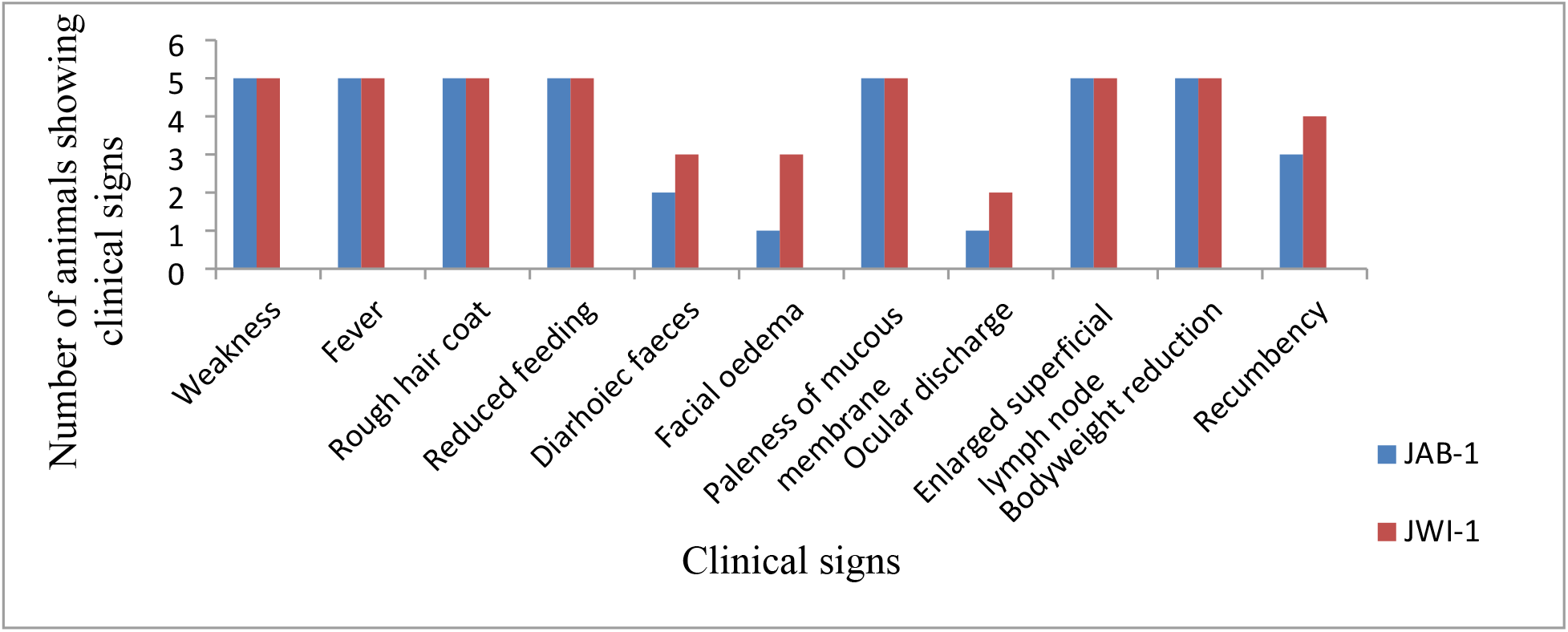
Different clinical signs observed during the study period in experimentally infected goats with *T*.*congolense* isolates from two tsetse infested (Jawi=JWI-1 and Jabitehenan= JBI-1) areas of northwest Ethiopia.

**Table 2:**
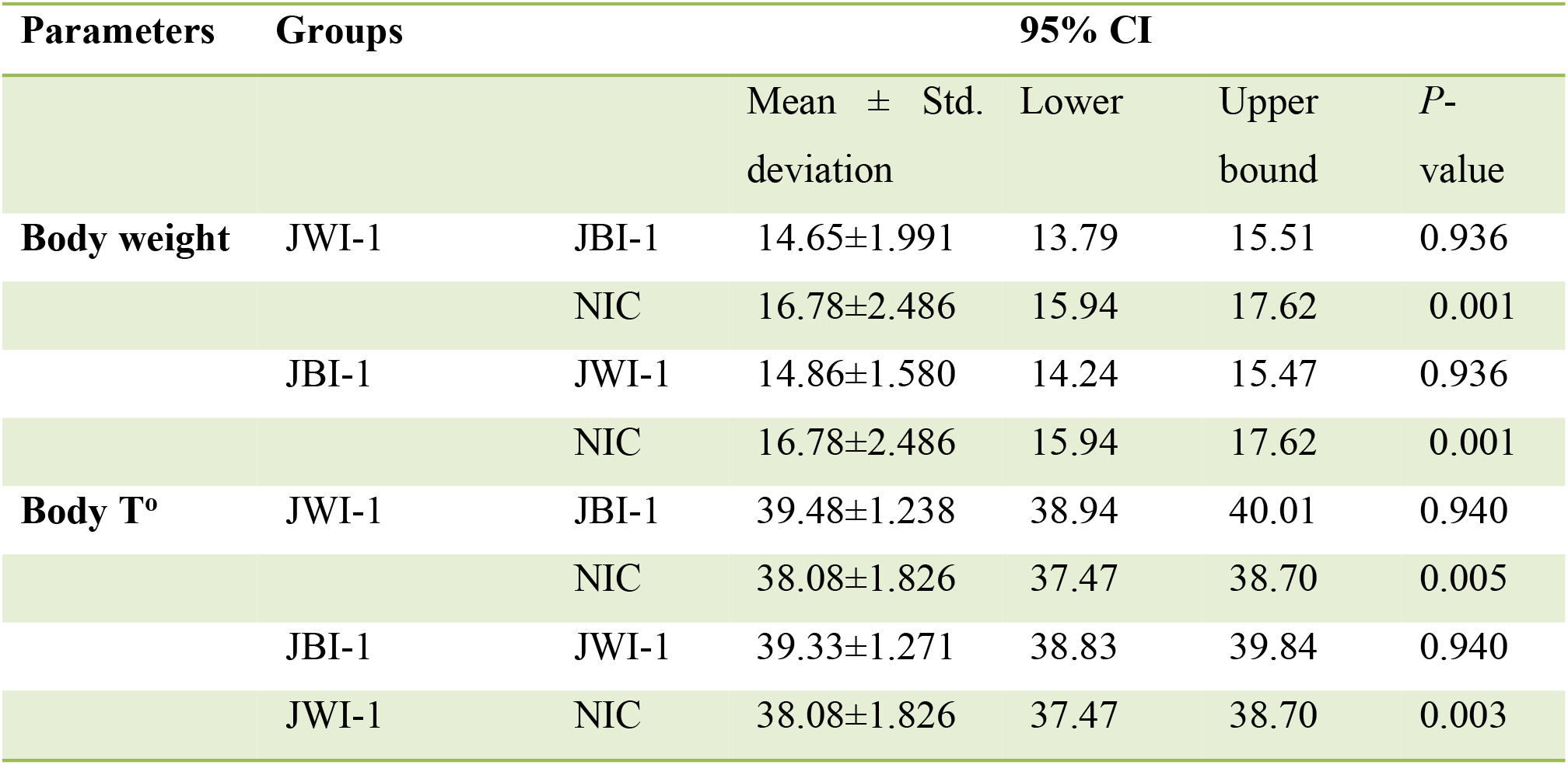
Multiple comparisons of mean body weight and body temperature between the trypanosome infected and none infected and as well as between the two areas isolates of *T. congolense* infected groups.

The mean rectal temperatures of infected groups (39.33±1.27°C, 39.48±1.24°C for Jawi and Jabitehenan isolates respectably) were significantly higher (*P*<0.05) than the control group (38.5 1°C ± 1.82). However, there was no significant (*P*>0.05) difference between mean rectal tempera tures of the two areas *T. congolense* isolate infected groups. Infected goats temperature started rising since day 6 of Pi, concurrent with the appearance of parasitaemia, and then fluctuated throughout the study period. The highest mean temperature recorded was 41.45 °C, from group JBI-1 on day 35 of Pi (Fig. 4).

**Figure 4:**
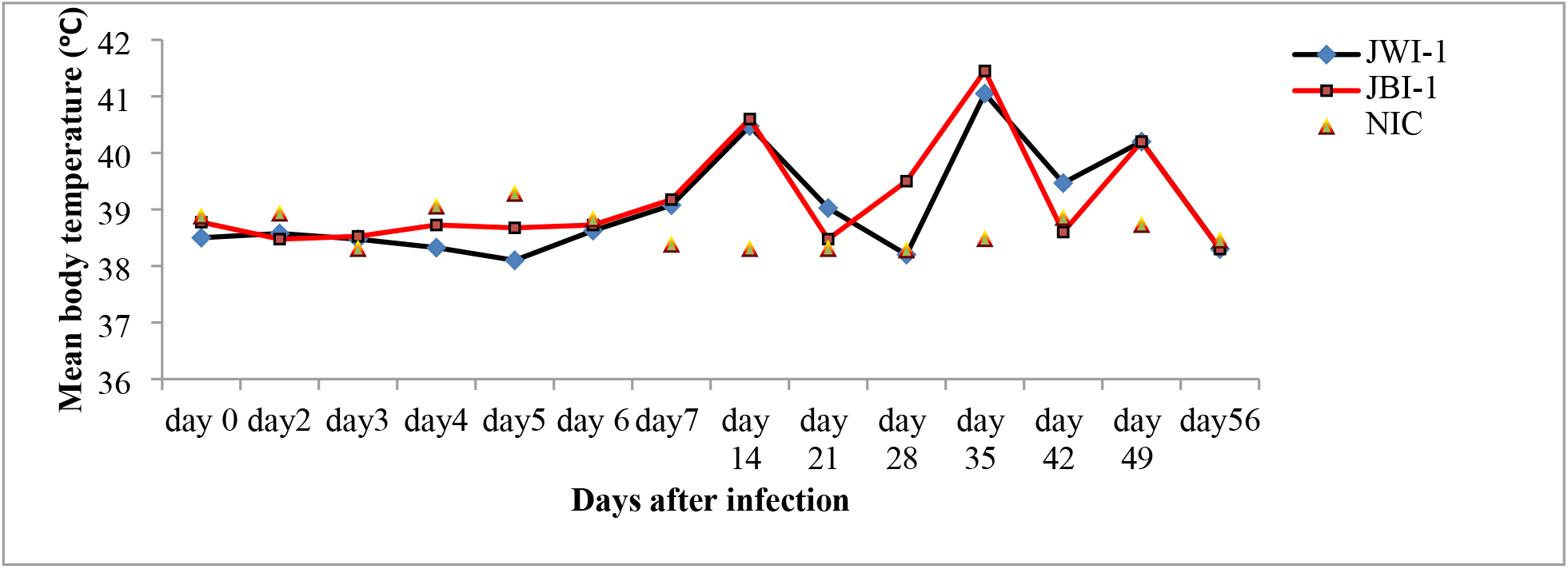
The mean body temperature measured during the study period in groups of non infected control (NIC) and experimentally infected with *T. congolense* isolates from two tsetse infested (Jawi=JWI-1 and Jabitehenan=JBI-1)areas of northwest Ethiopia.

As shown in figure (5), the infected groups significantly (*P*<0.05) lost body weight, while the control group steadily gained body weight. However, there was no significant body weight lost (*P*>0.05) difference between Jawi and Jabitehenan *T. congolense* isolate infected groups (Fig. 5).

**Figure 5:**
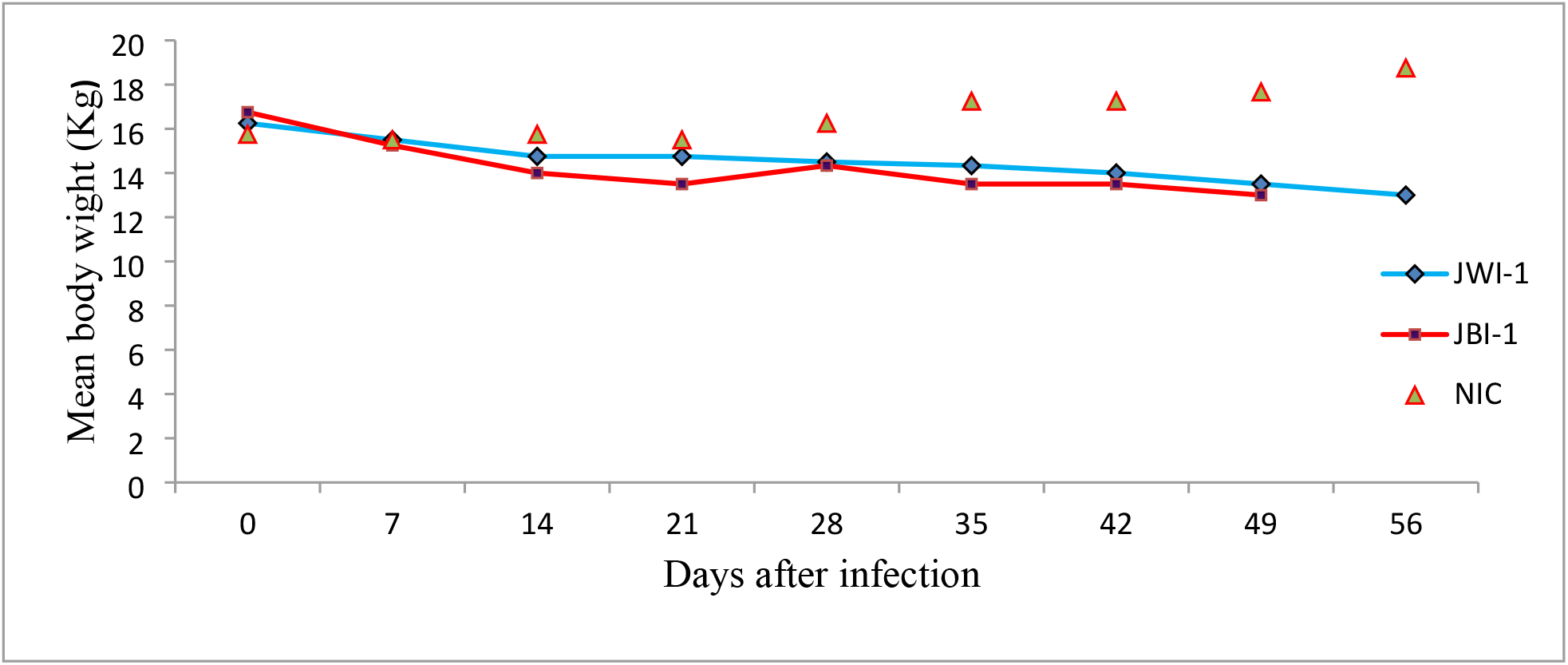
The Mean body weight measured during the study period in groups of non-infected control (NIC) and experimentally infected with *T. congolense* isolates from two tsetse infested areas(Jawi=JWI-1 and Jabitehenan=JBI-1) of northwest Ethiopia.

### 3.3. Hematological findings

The overall mean hematological changes observed in all experimental animals during the study period were summarized in Tables3. Significant reductions (*P*< 0.05) of mean PCV, Hgb concentration, total RBC and WBC counts were detected in all infected groups as compared with the non-infected control group. The difference between JWI-1 and JBI-1 mean PCV was also significant (*P<*0.05) that goats infected with Jabitehenan *T. congolense* isolates (JBI-1) showed low PCV record (Fig.6).

**Table 3:** Mean haematological values of experimentally infected goats with *T*.*congolense* isolates of Jawi and Jabitehenan areas

| Hematological values | Group | Mean ± SD | 95 % CI for mean | P- value |
| --- | --- | --- | --- | --- |
| PCV values (%) | Jawi isolate | 22.00± 3.86 | 20.50-23.50 | <0.05 |
| | Jabitehenan isolate | 18.43±5.759 | 15.94 -20.93 | |
| | Negative control | 27.31±1.19 | 26.90-27.71 | |
| Hgb concentration (g/dl) | Jawi isolate | 7.29±1.36 | 6.76-7.81 | <0.05 |
| | Jabitehenan isolate | 6.35±1.97 | 5.50-7.20 | |
|  | Negative control | 9.00±0.76 | 8.74-9.26 |  |
| **RBC count (×106/μl)** | Jawi isolate | 5.75 ± 0.91 | 5.42-6.08 | <0.05 |
|  | Jabitehenan isolate | 4.85 ± 1.20 | 5.42-6.08 |  |
|  | Negative control | 9.23±3.05 | 8.85-9.88 |  |
| **MCV (fl)** | Jawi isolate | 27.46±2.82 | 26.37-28.56 | <0.05 |
|  | Jabitehenan isolate | 27.39±3.99 | 25.67-29.11 |  |
|  | Negative control | 22.94±1.67 | 22.38-23.51 |  |
| **MCH (pg)** | Jawi isolate | 9.93±4.29 | 8.27-11.59 | >0.05 |
|  | Jabitehenan isolate | 8.91±1.276 | 8.77-16.45 |  |
|  | Negative control | 8.78±1.222 | 7.21-7.68 |  |
| **MCHC (g/dl)** | Jawi isolate | 33.33±1.992 | 32.57-34.09 | >0.05 |
|  | Jabitehenan isolate | 32.86±1.96 | 32.00-33.73 |  |
|  | Negative control | 32.47±1.61 | 31.93-33.02 |  |
| **WBC count (×103/μl)** | Jawi isolate | 7.17±2.12 | 6.36-7.98 | >0.05 |
|  | Jabitehenan isolate | 7.04±1.55 | 6.37-7.71 |  |
|  | Negative control | 7.91±1.79 | 7.53-8.29 |  |

**Figure 6:**
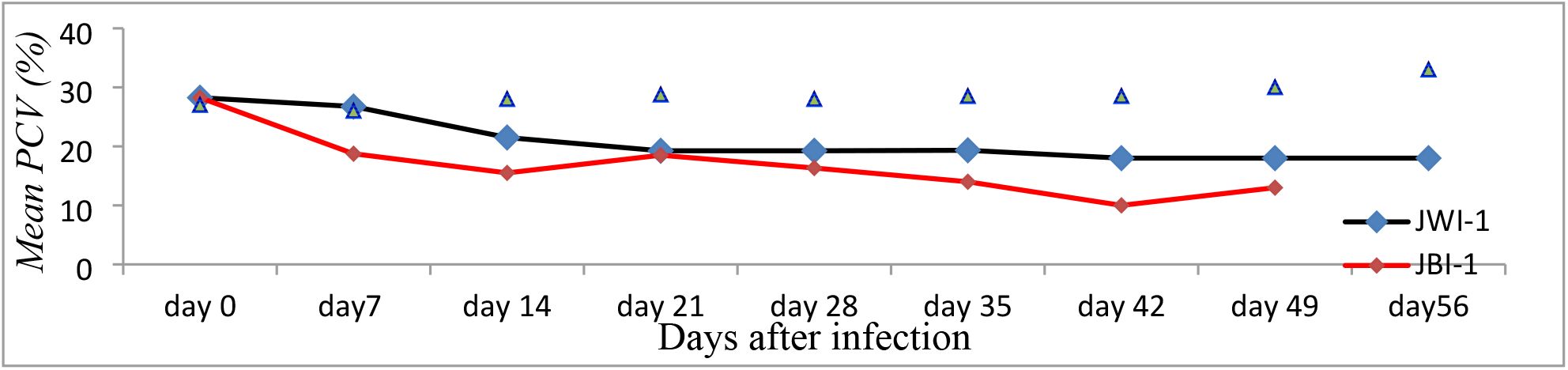
Mean PCV percentage in groups of non-infected control (NIC) and experimentally infected goats with *T. congolense* isolates from two different tsetse infested areas of northwest Ethiopia.

Infected groups had significant reduction (*P* <0.05) of mean Hgb concentration as compared with the non-infected control group (Fig. 7). Experimental infection with Jabitehenan *T. congolense* isolates (JBI-1) did have significantly (*P* <0.05) lower Hgb concentration as compare to those infected with *T*.*congolense* isolated from Jawi (JWI-1).

**Figure 7:**
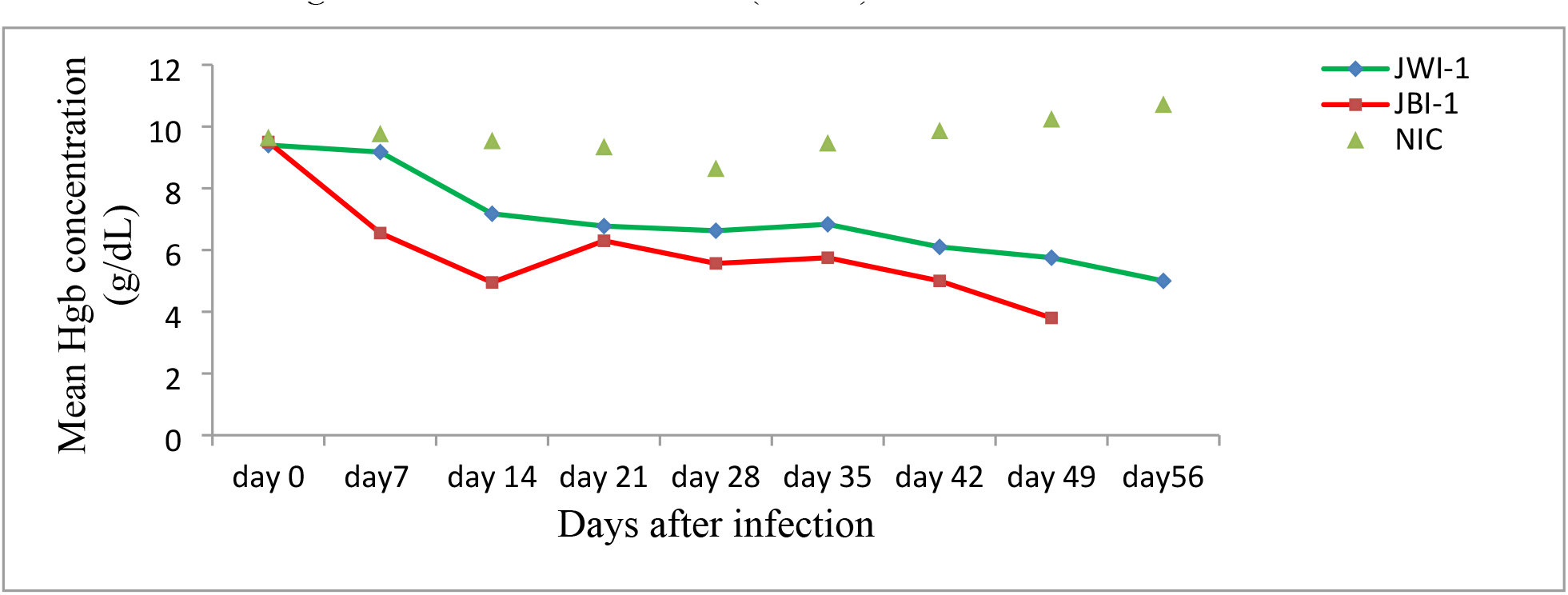
Mean Hgb concentration of non-infected control (NIC) and experimentally infected goats with *T. congolense* isolates from two different tsetse infested areas of northwest Ethiopia.

**Figure 8:**
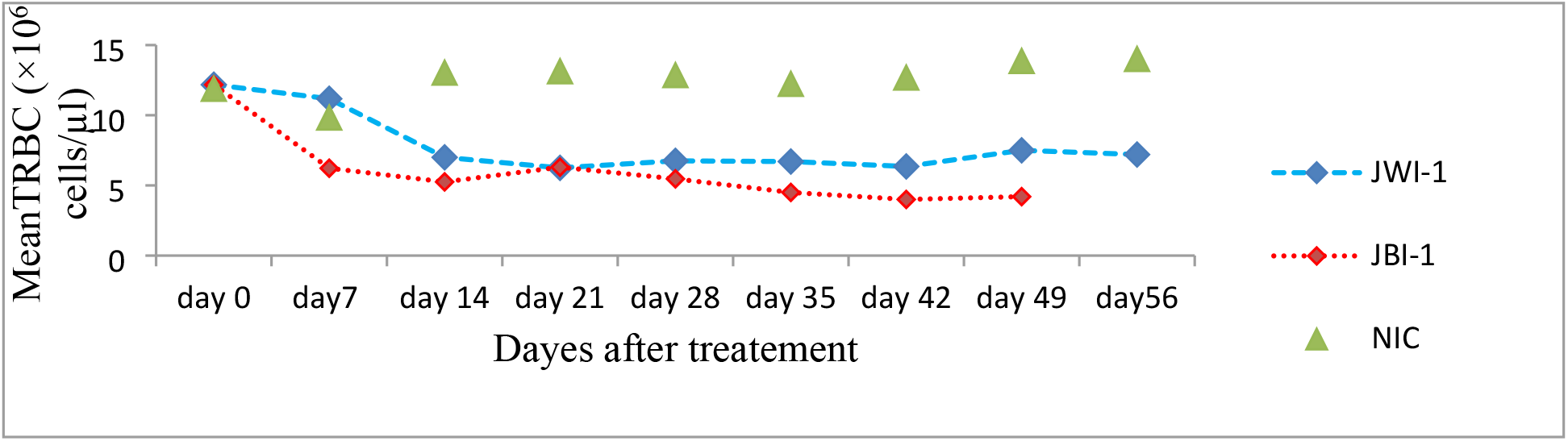
The Mean RBC count of non infected control (NIC) and experimentally infected goats with *T. congolense* isolates from two different tsetse infested areas of northwest Ethiopia.

Total RBC counts of infected groups were also significantly lower (*P* <0.05) than those of goats in none infected control groups. A significance difference (*P* <0.05) was also seen between JBI-1 and JWI-1 groups with the Jawi isolate infected groups showed higher RBC count (Fig. 7).

The mean total WBC count of infected groups was significantly lower (*P <*0.05) than the non infected control groups (Fig. 9). However, there was no significant (*P >*0.05) mean WBC count difference between infected groups.

**Figure 9:**
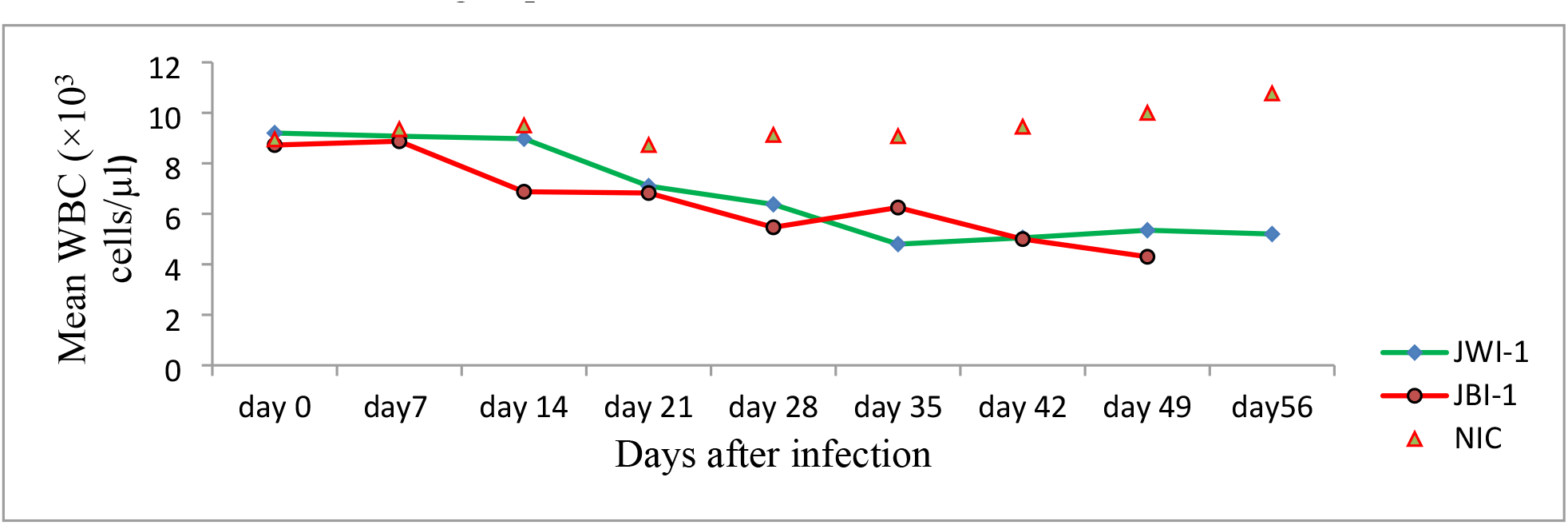
Mean WBC of non infected control (NIC) and experimentally infected goats with *T. congolense* isolates from two different tsetse infested areas of northwest Ethiopia.

Mean MCV value (**Fig. 10**) Showed a significant (*P* <0.05) difference between infected and none-infected (NIC) groups. But there was no significant (*P* >0.05) MCV value difference between the two areas *T*.*congolense* isolate infected groups. No significant difference (*P* >0.05) was also observed between mean MCH of infected groups and NIC groups. The mean MCH value was not significantly (*P* >0.05) different between the two areas *T*.*congolense* isolate infected groups. MCHC values (*P* >0.05) did not show significant difference between infected and non-infected groups. Furthermore, there was also not significant (*P* >0.05) variation of MCHC value between JWI-1 and JBI-1 groups.

**Figure 10:**
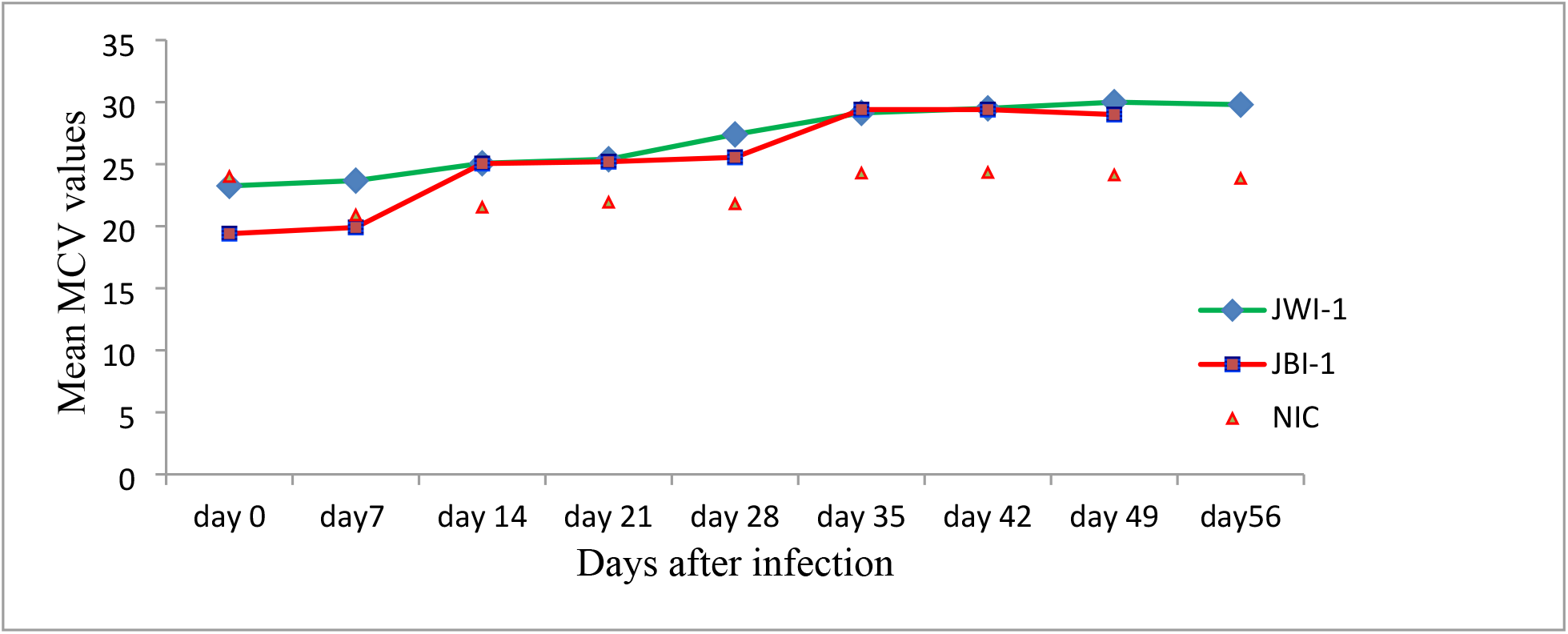
Mean MCV values of non infected control (NIC) and experimentally infected goats with *T. congolense* isolates of Jawi and Jabitehenan areas.

From the mean differential WBC counts value, neutrophils level were significantly (*P <*0.05) depressed in infected groups as compared to the non infected control groups and no significance (*P >*0.05) difference was observed between the two infected groups. Monocytes and eosinophil’s level showed insignificant increment in infected groups. However, Lymphocyte count showed insignificant decrement in infected groups. The basophiles count in all the three infected groups fluctuated within the range level of 0 and 1throughout the experiment and no significant (*P >*0.05) variation was detected among experiment groups based on the changes of numbers of basophiles during the experiment.

### 3.4. Postmortem findings

Postmortem examination was done carefully in each of the euthanized groups of experimental animals and the major findings include enlarged and edematous spleen; enlarged and congested liver; pale and swelled kidney; pneumonic lungs especially with rubbery consistency and meaty appearance and enlarged and congested heart. More accumulation of body cavity fluid than normal in the chest, lungs, abdomen and pericardium was also observed (Fig.11).

**Figure 11.**
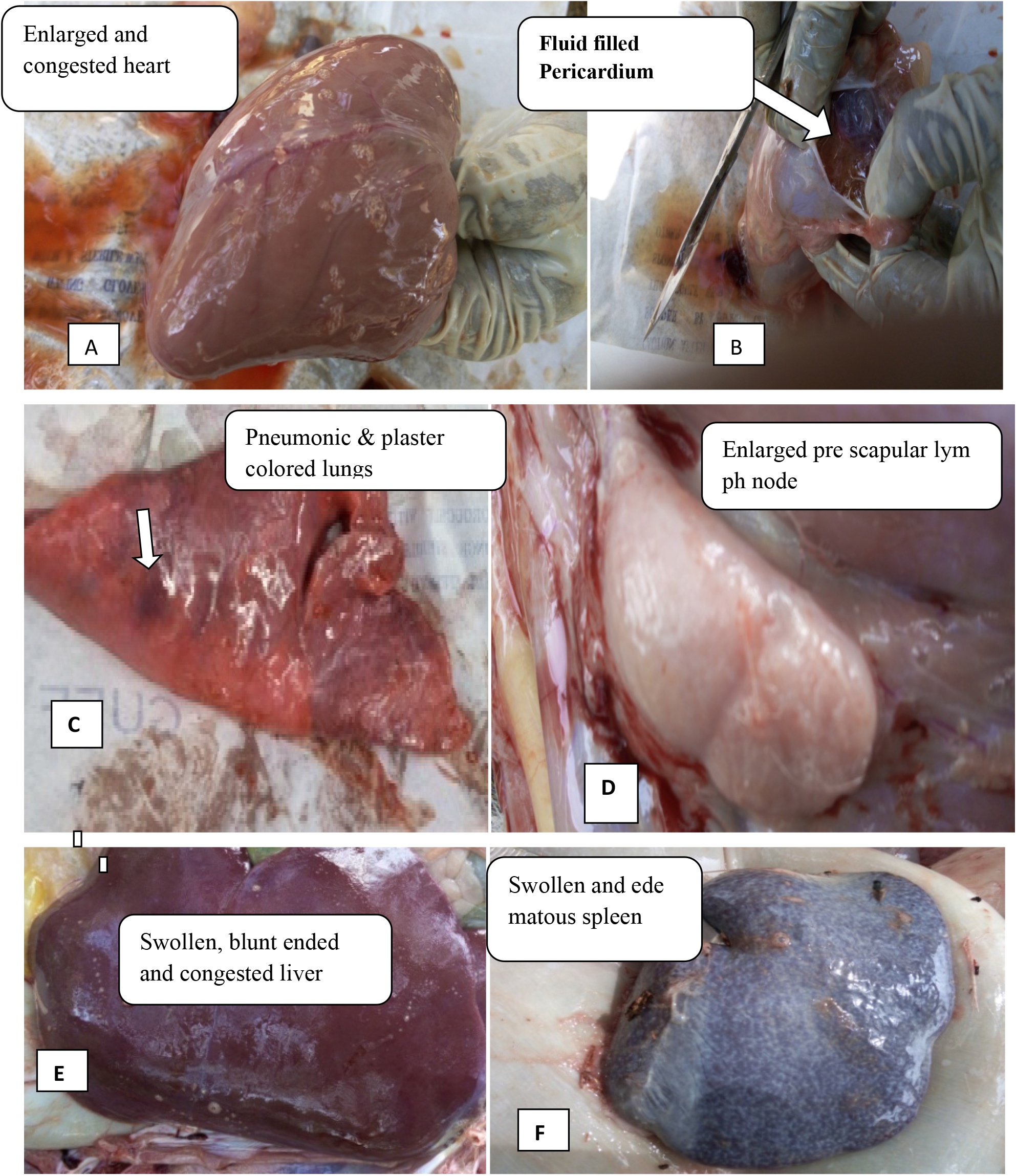
Major gross pathological findings in goats of experimentally infected with *T. Congolense* isolates of Jawi and Jabitehenan areas. **A**. Enlarged and congested heart from Group JBI-1;**B**. Excessive fluid in pericardium from Group JBI-1;**C**. pneumonic lungs especially with rubbery consistency and meaty appearance from group JWI 1; **D**. Enlarged pre scapular lymph node from JWI 1; **E**. Swollen, blunt and congested liver from group JWI-1; **F**. swollen and edematous spleen from Group JBI-1.

### 3.5. Drug sensitivity test

#### 3.5.1. Parasitaemia changes

Treatment was given on day 14 of pi according to their respective groups when experimental animals got peak parasitaemia. Both isometamidium chloride and diminazene aceturate treatment significantly reduced *T*.*congolense* parasitaemia from 24 hours of post treatment (pt). There was also reduction of the severity of clinical signs as compare to the infected control group. But latter *T*.*congolense* was observed in these treated animals’ buffy coat microscopy. The first recurrence was discovered in a goat in group 6 (JBI-ISM) on day 42. Another goat from group 6 (JBI-ISM) was also found to have parasitaemia on day 49 of pt. On day 51, relapsing parasitaemia was visible in a goats from group 3 (JWI-ISM) and two goats from group 2 (JBI-DA).

#### 3.5.2. PCV value change

The PCV improvement was tested by comparing the mean PCV value of before treatment and after treatment (Table 5). The mean PCV and body weight (bwt) improvement was calculated using the following formula (28).

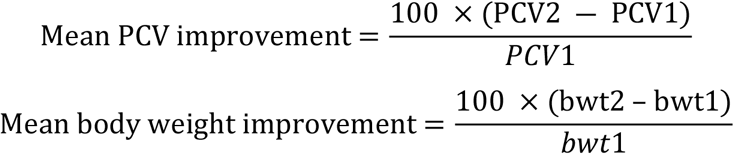

The PCV values of all four infected then treated groups of experimental animals were below the standardized normal caprine PCV value (22-38% by Meredith and Robin, (29)). Between day15 to day 58 of pt an improvement of mean PCV values were seen in all experimental animal groups except group JBI-DA. JWI-DA group showed higher mean PCV improvement (9.64%) during this period. However, the overall PCV value was still below the normal level. PCV reductions in all of the treated groups were also seen in the third phase of study (day 59-100). So trypanocidal treatment was unable to return the normal PCV values of infected animals’ effectively.

**Figure 13:**
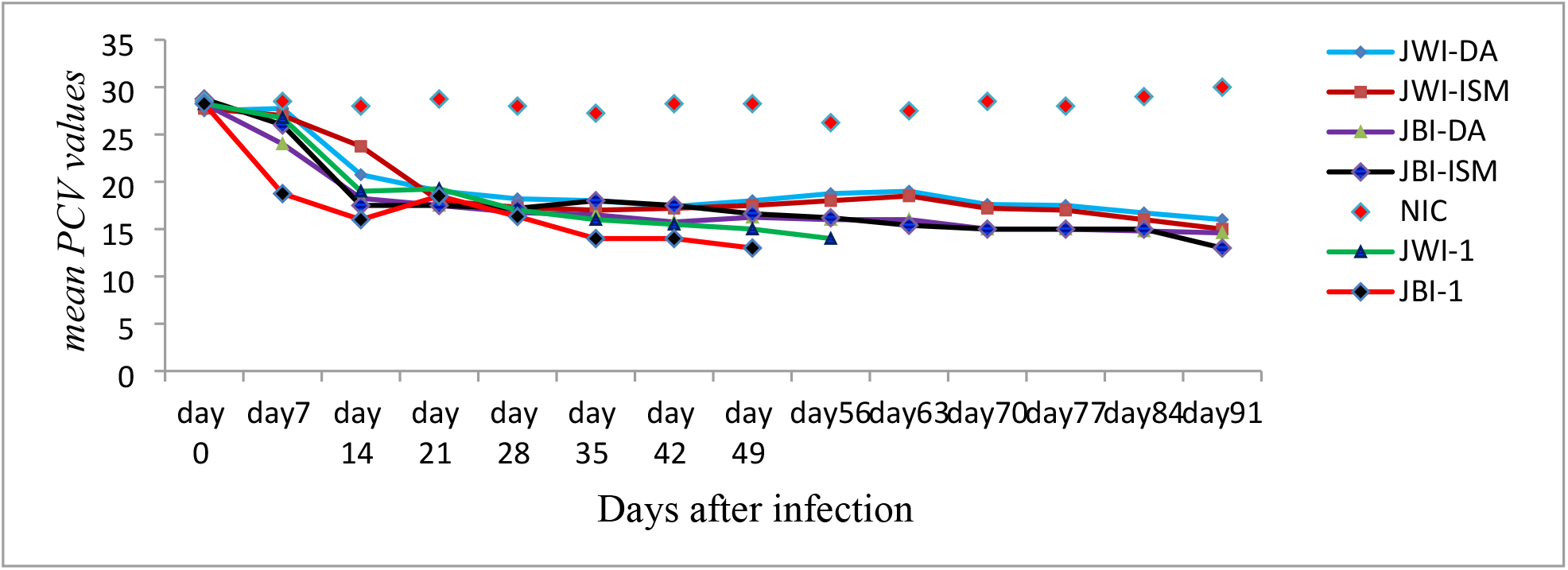
Mean PCV values of none infected (NIC), infected but not treated (JWI-1 and JBI-1) and treated (JWI-DA, JWI-ISM) groups of goats after experimentally infected with *T. congolense* isolates of Jabitehenan and Jawi

#### 3.5.3. Body weight change

JWI-DA group shows the highest body weight gain improvement (3.33%) and Group JBI-ISM was the least (−13.84).

The mean body weight measurement of negative control (NIC) group was significantly higher than all of the mean body weight of treatment groups (*P* <0.05) throughout the experiment period (Table 6). The mean body weight measurements observed in the study was not showed significant difference (*P*>0.05) up to day 21 pi between infected, infected then treated and negative control groups (fig.14).

**Figure 14:**
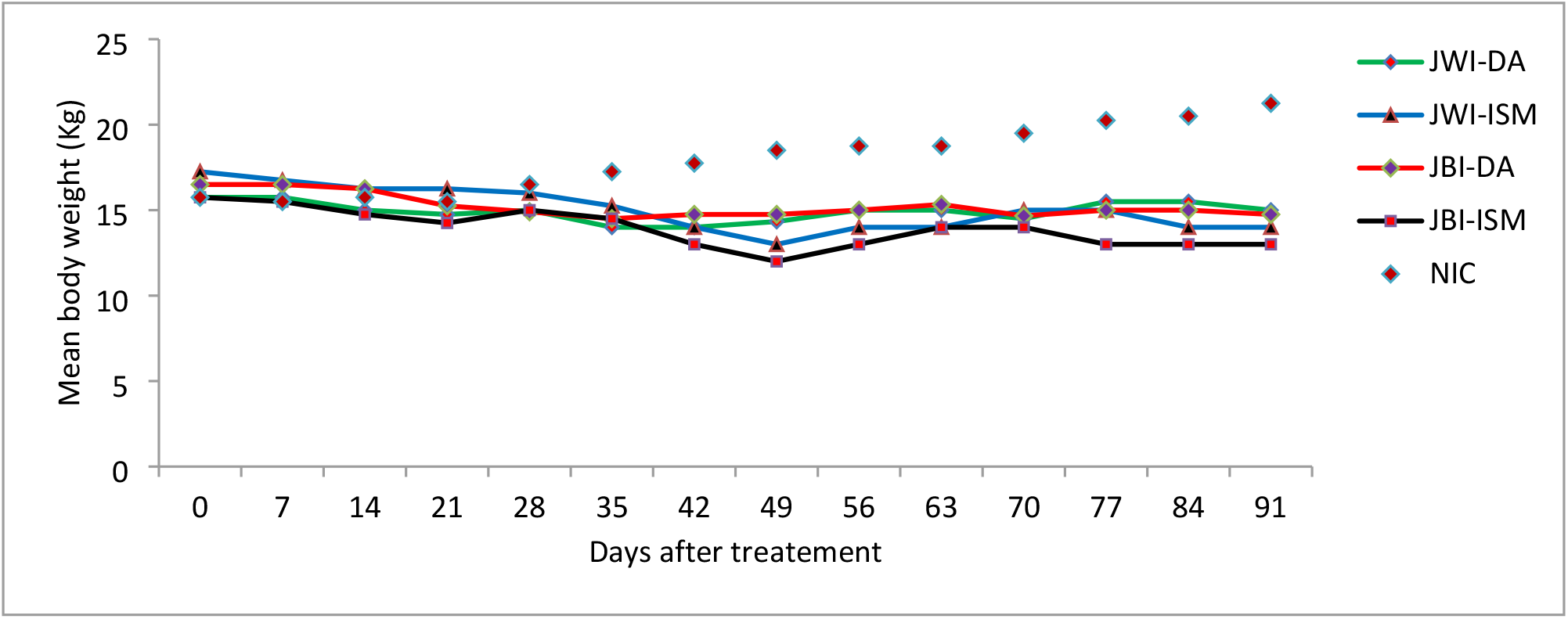
The mean body weight of none infected (NIC) and infected then treated (JWI-DA, JWI-ISM, JBI-DA, JBI-ISM) groups of goats after experimentally infected with *T. congolense* isolates of Jabitehenan and Jawi areas.

## 4. DISCUSSION

### 4.1. Development of parasitaemia

The first parasitaemia, in goats infected at field for transporting of the parasite to the experimental site was detected on day 11 and day 13 pi for Jabitehenan and Jawi isolates respectively and peak parasitaemia for the respective groups was also observed on day 26 and 29 of pi. This was long time as compare to the experimentally infected goats at the experimental site up on which parasitaemia was observed on day 5 of pi in Jabitehenan isolate infected groups and on day 6 of pi for Jawi isolates infected groups. The reasons for this parasitaemia detection time difference between the field infected and the experimental infected goats may be due to the fact that artificial transferring of the parasite between two different host animals species (from cattle to goats) may cause stress and instability of parasites to attain its proliferation capacity than transferring of parasites between the same host species of animals (from goat to goat). The other possibility may also be due to the reason that syringe transferring (artificial inoculation) for the field inoculated parasites was for the first time. However, the parasite that was inoculated to goats at the experimental site may adapt syringe transferring mechanisms for the second time than field inoculated parasites. The latter suggestions is supported by a research works on *T. brucei* (30) and *P. chabaudi* (31) that had shown syringe passage had effects for rapid increments of parasitaemia. Syringe-passaging is a laboratory technique that selects for virulent parasites, but these parasites don’t behave like naturally virulent stocks, according to Turner *et al*., (30).

Infection doses were similar for both types of parasites and the experimental animals were grouped randomly; managed similarly and sampling days were identical. Therefore, it is believed that the difference in time for the onset of parasitaemia in the two areas *T. congolense* isolates could be genetic basised.

Sewell and Brocklesby (32) affirmed in ruminants that a prepatent period of *T*.*congolense* is 3-5 day. Abenga *et al*. (33) and Ezeokonkwo *et al*. (34) from a trial on dog reported that the prepatent period of *T*.*congolense* in artificially infected dogs was 12 days. This prepatent period difference may be due to differences in specious of experimental animals and strains of isolates of the pathogen. Similarly Hoare, (35) reported that the prepatent periods of trypanosome is variable, depending on the host and the parasites isolates.

Since the initial infection, parasitaemia increased progressively to reach at peak level on day 14 of pi and continue as fluctuating wave. The observed fluctuating parasitaemia in the infected goats was similar to the trend seen by Ezeokonkwo *et al*., (36). Similarly Dagnachew *et al*. (13) confirmed that trypanosomosis is characterized by waves of parasitaemia.

High level of parasitaemia probably overwhelmed the immune response of goats within a short period of time. Hence, severity of infection in all infected groups within a short period of time (27-59 days pi) was occurred. In agreement with this finding, Ezeokonkwo *et al*. (34) observed that the survival period of dogs infected with *T. congolense* was 1-4 weeks. However, the chronic disease syndrome seen in ruminants characterized by low parasitaemia with many waves in which the infected animals survive for many months (37; 38) was contradictory to the infection seen in this study.

### 4.2. Clinical findings

Reduction of feed intake, weakness, fever, rough hair coat, enlarged superficial lymph nodes, Lacrimation, weight loss, paleness of mucus membranes, and finally recumbence were seen in all of the experimental goats. Manifestations of typical clinical signs of trypanosomosis in *T*.*congolense* infected calves: such as depression, fever, in appetence, swelling of pre-scapular and pre-femoral lymph nodes, rough hair coat, and overall reduction in mean PCV below 20% were reported by Hagos *et al*. (39).

Depression, dullness, ocular discharge, anemia, pyrexia and bottle jaw which had observed in thi s study were reported as characteristic of trypanosomosis in animals (40). The current finding was also similar with the finding of Biryomumaisho *et al*., (41) who study on *T*.*congolense* and *T. brucei* infection in Small East Africa goats. Ezeokonkwo *et al*. (34) on dog *T. congolense* infection reported that clinical signs like; pallor of the mucous membrane, depression, ocular discharge, anorexia and enlargement of the superficial lymph nodes were characterized. These clinical findings were similar to the findings by Dagnachew *et al*. (13) who analyzed and classified the early and late clinical findings in young zebu cattle infected with *T. vivax*. According to Seed and Hall (42), pyrexia is due to the metabolism of tryptophan to tryptophol by trypanosome infections. The accumulation of tryptophol in animals’ body is responsible for febrile conditions as a consequence of humoral antibody response to heterologous antigens. A significant incensement of temperature in *T*.*congolense* infected animals depends on the enhanced level of released pyrogens in the severely stressed animals (43).

Furthermore, loss of body weight during *T. congolense* infections in domestic animals has been reported (44) indicating the economic impact of the disease. Similar to Nwoha and Anen (43), loss of body weight in *T. congolense* infection may be resulted from mobilization of body energy reserves due to deprivation of essential nutrients for the synthesis of ATP in the anorexic goats. So the more pronounced loss in body weight of the infected group might be the sequel of being in anorexia. The cause of sub-mandibular oedema may be with similar possibilities reported by Pentreath, (45) that increased cellular trafficking (especially macrophages) across the capillary endothelium, accumulation of substances such as proteases which was released by the parasite or a range of substances released from tissues by the parasite (including vasoactive amines and was a sequence of hypoalbuminemia. Tissue barrier damage in the kidney glomeruli which allowing the pathological loss of a number of substances from serum and such that a cause for sub-mandibular and other lower body parts oedema was also reported by Pentreath and Kennedy (46).

### 4.3. Haematological findings

The pragmatic reduction of PCV, total RBC count and Hgb concentration following infection indicates anaemia which is a cardinal feature of trypanosomosis in animals (47; 48).The reduction in mean PCV values might have a direct correlation with the decline in total RBC counts. The finding of low PCV value was similar with the finding by Hagos *et al*. (39). Many factors have been reported in the literature to be responsible for the reductions in total RBC counts, Hgb concentrations and the subsequent PCV value (which communally are termed as anemia) in trypanosomosis of livestock. Red blood cell counts can be reduced as a result of increased erythrophagocytosis which was reported as an important mechanism leading to anaemia in the pathophysiology of *T. congolense* infection in Zambian goats (49).From these factors erythrophagocytosis (50), disorders of coagulation (51), increased plasma volume and haemodilution (50), dyshaemopoiesis in which the bone marrow fails to produce RBC and immune mediated haemolysis (52) were the common reasons in different literatures. Apart from these postulation Ezeokonkwo *et al*. (34) reasoned for this phenomenon as it is due to hyperspleenism in which the red blood cells become prematurely damaged by the prevailing injurious factors in the splenic environment. These phenomena may have played an important role in this study since the infection caused a marked enlargement of spleen in all experimentally infected goats.

Reduction in the total number of WBC (leucopenia) in the infected group implies that *T. congolense* had an immunosuppressive effect on these groups of goats leaving the goats with an impaired immune defensive mechanism; hence severity of all goats in that group within a short periods of time. The leucopenia recorded supported the long held view that trypanosome infection of livestock had an immunosuppressive effect on the infected animals. Immune depression in trypanosomosis is a well recognized and well studied characteristic of trypanosomosis in livestock, humans and mice (53).

According to Anosa *et al*. (48) trypanosomosis due to *T. congolense* leads to a reduced capacity to mount a primary humoral immune response to non-trypanosome antigens and an inferior response to vaccines. The end result is that the immune-compromised host may be less able to control the infecting trypanosome population, other concurrent diseases or less respond normally to vaccination regimes.

The phenomenon of leukopenia in the current study finding was also similar to the observations of Maxie *et al*. (54) in which pancytopenia, i.e. anaemia, leukopenia, and thrombocytopenia associated with *T. vivax* and *T. congolense* infections of cattle were described. However, this finding was disagreed with the report of Onyeyili and Anika (55) who observed leukocytosis in dogs and rats infected with *T. brucei*. The Leucopenia occurrence might be due to inappropriate responses or decreased stimulation of the bone marrow during trypanosomosis and/or immunosuppressive actions of trypanosome infection (56).

The severity of clinical signs and haematological value reduction were more in Jabitehenan isolate infected groups than Jawi isolate infected groups of experimental goats. Not only the severity of clinical signs and reduction of haematological values but also the time duration to develop these signs was short in Jabitehenan isolates. The earlier severity of infection among the infected groups of goats was recorded on day 27 post infection from Jabitehenan isolate infected groups. This clinical sign output gives a clue that there may be presence of inter-isolate pathogenicity variation between *T. congolense* isolates of Jawi and Jabitehenan.

The current study showed macrocytic normochromic type of anemia as it was seen that there was increment of the average size of each RBC (MCV) and unchanged MCH and MCHC values.MC V was found to be significantly increased while MCH, and MCHC, values did not differ significantly from those of none infected control groups and this signifying macrocytic normochromic RBC (57). Macrocytosis in African trypanosomosis usually arises from the circulation of large numbers of reticulocytes during the acute phase of the anaemia (58; 59). Slight and sporadic increases in the values of MCH were probably due to free circulating haemoglobin arising from hemolysis of RBC (60). MCHC was reported to be normal in *T. congolense* infected cattle (61), while it was depressed in *T. brucei* infection of rabbits (62). MCH was also unaltered in *T. brucei*-infected rabbits (62).

### 4.4. Postmortem findings

The major postmortem findings include enlarged and edematous spleen, enlarged and congested liver, pale and swelled kidney along with enlargements of heart. Petechial haemorrhage in the lung and enlarged lymph nodes were common findings. More than normal fluid accumulation in body cavities, oedema, (in the chest, lungs, abdomen and pericardium) was also observed.Similar pathological findings were reported by Morrison, (63). Since *T. congolense* is from haematinic group (64), it produce numerous changes in the cellular and biochemical constituents of blood (65).Damage to endothelial cells by parasite products, immune complexes, vasoactive amines and cytokines increases vascular permeability so that excess fluid accumulation (oedema) in the lower parts of infected animals body were occurred. In *T. congolense* infections, a generalized dilatation of capillary beds, which alters the haemodynamic, was observed and reported (66).The concomitant anaemia and more sluggish tissue perfusion affect the exchange of metabolites and are associated with intracellular oedema of capillary endothelial cells. Fibrinous microthrombi form in response to endothelial damage. Alterations to the microcirculation produce secondary degenerative changes in tissues. As capillary permeability increases, phagocytes and products of the parasite extravasates more readily and are responsible, in part, for some of the tissue lesions (63). Enlargements of most of reticulo-endothelial organs were common in all of the infected groups in the current study. Similarly, Orhue *et al*. (67) observed and stated that an important feature of the pathogenesis of trypanosomosis is the effect on lymphoid tissue. As the disease progresses, the volume of tissue in the spleen, lymph nodes and bone marrow increases markedly; so that enlargements of different organs were seen. This hyperplasia of reticulo-endothelial cells reduces lymphoid cell density, and eventual lymphoid depletion can occur (67).

### 4.5. Drug sensitivity tests

Both of the DA and ISM trypanocides were unable to completely treat trypanosomosis by *T. congolense* ever last in experimentally infected goats. Rather, relapsing of the parasitaemia was detected since twenty eight days of post trypanocidal treatment (Table-4). This could be taken as evidence of drug resistance in this stock of *T. congolense* contrary to the report by Fairclough(68)who reported that it was difficult to induce resistance to isometamidium even by repeated low dosages (0.25 - 0.5 mg/kg).

**Table 4:** Parasitaemia relapsing percentage of experimentally infected goats with *T*.*congolense* isolates of Jawi and Jabitehenan areas and then treated with DA and ISM

| Isolates origin | Treatment | Group | Relapse number | % |
| --- | --- | --- | --- | --- |
| Jawi | DA | (JWI-DA) | 0/5 | 0 |
| Jawi | IM | (JWI-ISM) | 1/5 | 20 |
| Jabitehenan | DA | (JBI-DA) | 1/5 | 20 |
| Jabitehenan | IM | (JBI-ISM) | 2/5 | 40 |

**Table 5:**
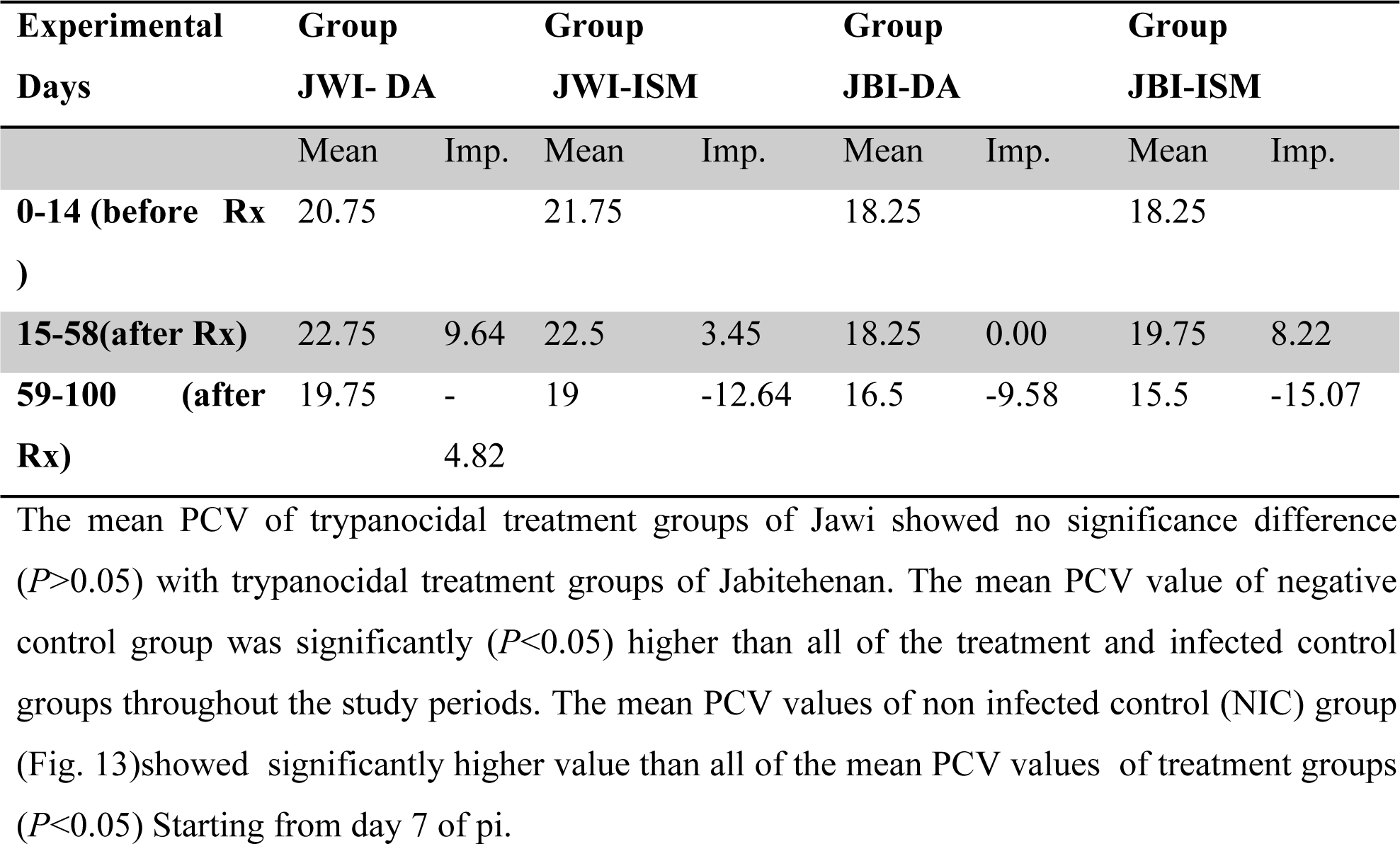
Mean PCV improvement of the infected then treated groups of experimental animals over the entire study period.

**Table 6:** Mean body weight improvement of the infected then treated groups of experimental animals over the entire study period.

| Experimental Days | Group JWI-DA |  | Group JWI-ISM |  | Group JBI-DA |  | Group JBI-ISM |  |
| --- | --- | --- | --- | --- | --- | --- | --- | --- |
|  | Mean | Imp. | Mean | Imp. | Mean | Imp. | Mean | Imp. |
| 0-14 (before Rx) | 15 |  | 14.75 |  | 16.25 |  | 16.25 |  |
| 15-58(after Rx) | 15 | 0.00 | 13 | -11.86 | 15 | -7.69 | 15 | -7.69 |
| 59-100 (after Rx) | 15.5 | 3.33 | 14 | -5.08 | 15 | -7.69 | 14 | -13.84 |

Imp= body weight improvement

This result was in agreement with Mulugeta *et al*. (69) who reported on the experimental study as subsequent to treatment with DA at a dose of 7.0 mg/kg bwt, trypanosomes reappeared in all 10 calves within 5-16 days and 0.5 mg/kg ISM trypanosomes reappeared in all animals within 4– 26 days post treatment. Such phenomenon clearly indicated the occurrence of resistant strains of *T. congolense* against either of the trypanocidal drugs (DA and ISM) treatment.This finding was also similar with Chaka and Abebe, (19) who reported on their experimental sensitivity trials conducted on mice using isolates of *T. congolense* taken from Ghibe, Bedele, Sodo and Arbaminch, and both IM and DA failed complete clearance of the parasite from experimental animal blood.

In agreement with this finding, Afework *et al*. (18) confirmed multiple drug resistance in cloned *T. congolense* isolated from Metekel region, northwest Ethiopia. According to Assefa *et al*. (15) Isometamidium chloride (ISM) at a dose of 0.5–4 mg/kg bwt and diminazine aceturate (DA) at a dose of 3.5–28 mg/kg bwt failed completely to cure *T. congolense* infections in mice. Similar resistance problems for *T. vivax* isolates against these drugs at higher dose were also reported by Dagnachew *et al*. (13). In contrary to this, Peregrine *et al*., (70) in Kenya also found 15 steers treated with 0.5 mg/kg isometamidium chloride were all cured. Another work conducted with two clones of *T. congolense* (from Uganda and Tanzania) showed them to be sensitive to the therapeutic activity of isometamidium chloride at 0.5 mg/kg.

The slight improvement in PCV readings after treatment (days 15-58pi) may be due to elimination of the drug sensitive population of trypanosomes from the animals’ body. However, the overall mean PCV is below the physiological value. This may be due to the presence of drug resistant populations. Body weight improvement in Group JWI-DA could be due to the probable effectiveness of DA in this group. The loss of weight in-Group 3, 5 and 6 may be attributed to the number of relapses experienced to trypanocidal drugs.

## 5. CONCLUSION AND RECOMMENDATIONS

All the infected goats showed an acute form of *T. congolense* infection accompanied by severe clinical signs. The Jabitehenan *T. congolense isolates that were* infected (JBI-1) groups showed more severe reductions in PCV, total RBC, total WBC, and Hgb concentrations. Furthermore, the severity of clinical signs in all infected goats by Jabitehenan T. congolense isolates occurred quickly in comparison to the Jawi T. congolense isolates infected groups (JWI-1). Therefore; this result gives a clue that the Jabitehenan isolate of *T. congolense* infection causes more severe disease and lethality within a short period of time. There was no satisfactory body weight improvement in all of the infected then treated groups. The PCV improvement was also insignificant throughout the study period. In addition, relapsing of parasitaemia was detected since day 28pt. This showed that the parasite develops drug resistant to both DA and ISM.

Therefore, based on the above conclusion the following recommendations were forwarded:

✤ Identification of T. congolence at molecular level was a great shortcome of this study. so further molecular characterization of *T*.*congolense* isolates should be done to know which kinds of sub-species are there in the region
✤ Trials to upgrade the efficancy of the present drugs and /or manufacture new anti-trypanosome drugs should be prompted.
✤ Implementation of integrated trypanosome control strategies is necessary to limit the problem in tsetse infested areas of northwest Ethiopia.

## 6. ACKNOWLEDGEMENTS

The authors acknowledge the University of Gondar, as this study was conducted under a megaproject sponsored by it. The College of Veterinary Medicine and Animal Sciences of the University of Gondar deserves acknowledgement as it permits different laboratories and equipment during the study. A sincere appreciation also goes to Bahir Dar Regional Veterinary Diagnostic and Investigation Center staff members for their absolute provision of logistic facilities and devotion of time in assisting all through the isolation of field parasites. The teaching farm staff at the University of Gondar are grateful for their permission to use experimental houses and their assistance in managing laboratory animals.

## Notes

### Competing Interest Statement

The authors have declared no competing interest.

